# Serine Deamination as a New Acid Resistance Mechanism in *Escherichia coli*

**DOI:** 10.1101/2022.05.05.490856

**Authors:** Michelle A. Wiebe, John R. Brannon, Bradley D. Steiner, Adebisi Bamidele, Alexandra C. Schrimpe-Rutledge, Simona G. Codreanu, Stacy D. Sherrod, John A. McLean, Maria Hadjifrangiskou

## Abstract

*Escherichia coli* associates with humans early in life and can occupy several body niches either as a commensal in the gut and vagina, or as a pathogen in the urinary tract. As such, *E. coli* has an arsenal of acid response mechanisms that allow it to withstand the different levels of acid stress encountered within and outside the host. Here, we report the discovery of an additional acid response mechanism that involves the deamination of L-serine to pyruvate by the conserved L-serine deaminases SdaA and SdaB. L-serine is the first amino acid to be imported in *E. coli* during growth in laboratory media, as the culture senesces. However, there remains a lack in knowledge as to why L-serine is preferred and how it is utilized. We show that in acidified media, L-serine is brought into the cell via the SdaC transporter and deletion of both SdaA and SdaB renders *E. coli* susceptible to acid stress, with a phenotype similar to other acid stress deletion mutants. We also show that the pyruvate produced by L-serine de-amination activates the pyruvate sensor BtsS, which in concert with the non-cognate response regulator YpdB upregulates the putative transporter YhjX, similar to what has been reported for this system during transition of *E. coli* to stationary phase. Based on these observations, we propose that L-serine deamination constitutes another acid response mechanism *in E. coli* that may function to protect *E. coli* as it transitions to stationary phase of growth.

**IMPORTANCE:** The observation that L-serine uptake occurs as an *E. coli* culture senesces is well-established, yet the benefit *E. coli* garners from this uptake remains unclear. Here, we report a novel acid resistance mechanism, where L-serine is deaminated to pyruvate and ammonia, promoting acid tolerance in *E. coli*. This study is important as it provides evidence of the use of L-serine as an acid response strategy, not previously reported for *E. coli*.

## INTRODUCTION

Acid stress is a substantial challenge to bacterial life. Acidic conditions can damage the bacterial cell envelope, rendering membrane-embedded proteins and the proton motive force established across the membrane dysfunctional or non-functional (1). Acidic conditions in the bacterial cell interior disturb vital physiological processes, such as enzymatic activity, protein folding, membrane- and DNA maintenance, all of which are needed for cellular function and can cause bacterial death. As a result, bacteria are equipped to withstand acidic conditions. One of the model bacterial organisms, *Escherichia coli*, occupies numerous environmental and host niches and encounters a range of acidic conditions in a niche-dependent manner. For example, all *E. coli* harbored in the gut are thought to be acquired via ingestion (2), indicating that *E. coli* must be able to survive the low pH of the stomach, which ranges from pH 1.5-3.5 (3). Uropathogenic *E. coli* (UPEC) strains – that colonize the gut asymptomatically for long periods of time (4) – oftentimes persist within the acidic environment of the vagina (pH 3.8-5) (5, 6) before ascending to and colonizing the urinary tract (pH 5.5-7) (7, 8). It is therefore not surprising that five acid resistance (AR) mechanisms, termed AR1-AR5, have been identified in *E. coli*, all of which have been shown to be active in the gut (9, 10).

The most well characterized AR mechanisms, AR2-5, depend on the import and subsequent decarboxylation of specific amino acids. The decarboxylation reaction consumes one proton, thereby increasing the cytoplasmic pH. AR2, which has been shown to be the most effective AR mechanism in *E. coli*, involves the deamination of glutamine to glutamate, which is then decarboxylated to form γ-amino butyric acid (GABA) (3, 11-13). AR3 involves the decarboxylation of arginine to produce agmatine, while AR4 leads to decarboxylation of lysine to produce cadaverine. Finally, ornithine decarboxylation to putrescine is the basis of AR5 (13). The AR1 mechanism of action is not well understood.

While decarboxylation of amino acids appears to be the primary means of acid resistance in *E. coli*, there is evidence that deamination of amino acids can also serve to increase the intracellular pH. This is evident in AR2, where glutamine deamination not only provides the glutamate that is then decarboxylated (AR2_Q) (14, 15), but also produces ammonia which consumes a proton and diffuses out of the cell as ammonium (14, 16). Here we demonstrate that the deamination of L-serine is an additional acid response mechanism in *E. coli*.

Notably, L-serine is the first amino acid consumed by *E. coli* when grown in complex media (17). Under aerobic conditions, it is known that L-serine is deaminated by SdaA and SdaB to produce pyruvate and ammonia (18). However, to date no metabolic role for serine deamination in *E. coli* has been described (19), as most of the pyruvate derived carbon is subsequently excreted from the cell (17). In *Klebsiella aerogenes* and *Streptococcus pyogenes* serine deamination has been shown to be important for regulating nitrogen balance and maintaining extracellular pH (20, 21) but the full import of serine deamination in these pathogens remains unclear. Given the lack of clearly defined purpose for serine deamination in *E. coli* as and the fact that at least one serine deaminase is expressed at all times (19), we hypothesized that serine may play a role, in mediating the acid response in *E. coli*.

Previous work indicated that L-serine induces the activation of the BtsS pyruvate sensor kinase (22), presumably due to production of pyruvate via SdaA/B mediated L-serine de-amination. Natural induction of *yhjX* has been previously reported by the Jung group to occur in *E coli* culture during mid-logarithmic growth phase (23), when bacterial cell density is high and nutrient depletion of LB begins to occur. We have previously shown that BtsS interacts with the non-cognate YpdB transcription factor to induce the upregulation of *yhjX* (23-25), a gene that is also induced under acidic conditions (26). In this work, we tested the hypothesis that L-serine deamination as a previously unrecognized acid resistance mechanism in *E. coli*. We show that under acidic conditions, serine is transported into the cell by the SdaC transporter and deaminated by the L-serine deaminases SdaA and SdaB, a process that imparts *E. coli* with protection from acid. We also demonstrate that the pyruvate produced by serine deamination is sufficient to induce *yhjX* transcription in a BtsS-YpdB dependent manner. Deletion of *sdaA and sdaB* leads to decreased acid resistance in *E. coli*. Collectively, our data uncover a previously uncharacterized acid resistance mechanism in *E. coli*.

## Methods

### Bacterial Strains and Growth Conditions

All studies were performed in uropathogenic *Escherichia coli* (UPEC) cystitis isolate UTI89 (27) and derived isogenic deletion mutants. For all analyses, strains were propagated from a single colony in lysogeny broth (LB) (Fisher Scientific), at pH 7.4 overnight at 37°C with shaking unless otherwise noted. Strains containing the luciferase reporter plasmid were grown in LB + 50 µg/ml gentamicin at 37°C, 220 rpm overnight. pH was measured using Thermo Scientific Orion Star A211 pH meter with an Orion™ Green pH Combination Electrode. Specific growth conditions for reporter and survival assays are described in the relevant sections below. Gene deletions were created using the λ-red recombinase system (28). The Δ*btsS*Δ*ypdB* and Δ*yhjX* mutant strains were created in previous studies (24). The *yhjX::lux* reporter, which was previously constructed (23), was introduced into each strain via electroporation and validated by PCR.

A complete list of primers, strains, and plasmids used for in this study can be found in Tables S1-S3.

### Luciferase reporter assay

Overnight cultures were spun at 3220 x g for 10 minutes. Pellets were resuspended in 5 ml 1X PBS and re-pelleted. Pellets were resuspended in 5 ml 1X PBS and normalized to a starting OD_600_ = 0.05 in 1 ml of the indicated media (LB, LB + 10 mM HCl, or LB + 50 mM MOPS or HEPES buffer + 10 mM HCl). Each suspension was used to seed black, clear bottomed 96 well plates at 200 µl per well from and grown at 37°C with shaking. OD_600_ and luciferase readings were taken every hour for 8 hours using a Molecular Devices SpectraMax i3 plate reader. At least 3 biological replicates were assayed.

### Acid Resistance Assays

Acid resistance assays were performed as follows: Overnight cultures were diluted 1:100 in 5 ml of LB and incubated at 37°C with shaking until they reached mid exponential growth phase. At that point, 1 ml of the culture was centrifuged at 16,000 x g for 5 minutes, washed in 1X PBS, serially diluted and spot-plated for CFUs. The pH of the remaining 4-ml culture was adjusted to a pH of 3 using 5 M HCl, and these cultures continued to incubate at 37°C, with shaking for an additional 30 minutes. A 1-ml sample was centrifuged, washed in PBS, and plated for CFUs. Percent survival was calculated by dividing the CFU/ml after acid stress by the CFU/ml of the input sample (collected before acid was added to the media).

### RNA Extraction and RT-qPCR

*Growth conditions for RNA sample collection:* To collect samples for transcriptional analysis, strains were grown aerobically in LB to an OD_600_ = 0.5 and then split into two conditions: continued growth in LB alone, or in LB in which HCl was added to a final concentration of 10 mM. Samples were taken for RNA extraction at time = 0, 15, 30, 60, and 120 minutes after splitting the culture. All samples were centrifuged at 6000 x g for 7 minutes at 4°C. The supernatant fractions were decanted, and cell pellets were flash frozen in dry ice and ethanol and stored at -80°C until RNA extraction.

RNA was extracted using the RNeasy mini kit from Qiagen, following the manufacturer’s extraction protocol. A total of 3 µg of RNA was DNAse treated using 2 units of Turbo DNase I enzyme (Invitrogen). A total of 1 µg of DNAse-treated RNA was reverse transcribed using Superscript III reverse transcriptase (Invitrogen/Thermo Fisher). cDNA was amplified in an Applied Biosystems StepOne Plus Real-Time instrument using TaqMan MGB chemistry with primers and probes listed in Table S1. All reactions were performed in triplicate with four different cDNA concentrations (100, 50, 25, or 12.5 ng per reaction). Relative fold difference in transcript abundance was determined using the ΔΔC_T_ method of Pfaffl *et al*., with a PCR efficiency of >95% (29). Transcripts were normalized to *gyrB* abundance. At least 3 biological replicates were performed for each transcript.

### Metabolomics

UTI89, Δ*btsS*Δ*ypdB*, and Δ*sdaC* were grown in LB and incubated at 37°C with shaking, until cultures reached an OD_600_ = 0.5. Then, 1 M HCl was added to the culture to a final concentration of 10 mM (pH=5). Cultures were incubated for another 15 minutes and then 1 ml of culture was collected. Cells were pelleted and supernatant was flash frozen and stored at -80°C until analyzed via Liquid Chromatography-Mass Spectrometry (LC-MS)-based metabolomics in the Vanderbilt Center for Innovative Technology (CIT). Isotopically labeled phenylalanine-D8 and biotin-D2 were added to 200 µL of culture supernatant per sample, and protein was precipitated by addition of 800 µL of ice-cold methanol followed by overnight incubation at -80°C. Precipitated proteins were pelleted by centrifugation (15,000 rpm, 15 min), and supernatants were dried down *in vacuo* and stored at -80°C. Individual samples were reconstituted in 120 µL of reconstitution buffer (acetonitrile/water, 90:10, v/v) containing tryptophan-D3, pyruvate-C13, valine-D8, and inosine-4N15. A quality control (QC) sample was prepared by pooling equal volumes from each individual sample. Quality control samples were used for column conditioning, retention time alignment and to assess mass spectrometry instrument reproducibility throughout the sample set and for individual batch acceptance.

LC-MS and LC-MS/MS analyses were performed on a high-resolution Q-Exactive HF hybrid quadrupole-Orbitrap mass spectrometer (Thermo Fisher Scientific, Bremen, Germany) equipped with a Vanquish UHPLC binary system and autosampler (Thermo Fisher Scientific, Germany). Metabolite extracts were separated on ACQUITY UPLC BEH Amide HILIC 1.7μm, 2.1 × 100 mm column (Waters Corporation, Milford, MA) held at 30°C. Liquid chromatography was performed at a 200 μL min−1 using solvent A (5 mM Ammonium formate in 90% water, 10% acetonitrile and 0.1% formic acid) and solvent B (5 mM Ammonium formate in 90% acetonitrile, 10% water and 0.1% formic acid) with a gradient length of 30 min.

Full MS analyses (6µL injection volume) were acquired over 70-1050 mass-to-charge ratio (m/z) in negative ion mode. Full mass scan was acquired at 120K resolution with a scan rate of 3.5 Hz, automatic gain control (AGC) target of 10e6, and maximum ion injection time of 100 ms. MS/MS spectra were collected at 15K resolution, AGC target of 2e5 ions, and maximum ion injection time of 100 ms. The acquired raw data were imported, processed, normalized and reviewed using Progenesis QI v.3.0 (Non-linear Dynamics, Newcastle, UK). All MS and MS/MS sample runs were aligned against a QC (pooled) reference run. Unique ions (retention time and m/z pairs) were de-adducted and de-isotoped to generate unique “features” (retention time and m/z pairs). Data were normalized to all features using Progenesis QI. Experimental data for measured serine (i.e., retention time and MS^2^ fragmentation pattern) and pyruvate (i.e., retention time) was consistent with reference standards.

### Statistical analysis

Statistical analyses were performed in GraphPad Prism. Details of sample size, test used, error bars, and statistical significance cutoffs are presented in the text or figure legends. All experiments were performed in at least three biological replicates. Representative graphs are shown for the luminescence reporter assays. qPCR data were analyzed using 2-way ANOVA with a Sidak’s multiple comparison test to compare individual time points. LC-MS abundance values were plotted as ArcSinh normalized values.

### Graphics

All graphical models and drawings were generated using BioRender.com

## Results

### *YhjX* is upregulated in response to low pH in a manner that depends on the non-cognate two-component system BtsS and YpdB

Previous studies have shown that *yhjX* is among the genes upregulated in response to acidic pH in K-12 *E. coli* (26). To confirm that these previous observations hold true in the UTI89 *E. coli* strain selected for our studies, we utilized a transcriptional reporter to monitor *yhjX* promoter activity under acidic conditions. We used a previously constructed strain UTI89/P*yhjX*::lux(24) that harbors a plasmid containing the *yhjX* promoter fused to the *luxCDABE* operon (23). Luminescence and bacterial growth were monitored over time in lysogeny broth (LB) with shaking. Cultures grown in neutral pH (7.4) showed a peak in *yhjX* promoter activity at 180 minutes, coincident with late logarithmic phase of growth (Fig. 1b, Fig. S1, and Supplementary file 1). Addition of increasing concentrations of HCl to the media led to an increase in *yhjX* promoter activity that was proportional to the drop in pH (Fig. 1b and Supplementary file 1). The addition of HCl in the conditions tested did not affect bacterial growth (Fig. S1). Addition of either MOPS or HEPEs buffers to the acidified culture media restored neutral pH and suppressed *yhjX* promoter activity to levels observed when grown in LB with pH 7.4 (Fig 1c and Supplementary file 1). These results indicate that *yhjX* is indeed an acid-induced target and corroborate previous observations in K-12 strains of *E. coli*.

**Figure 1.**
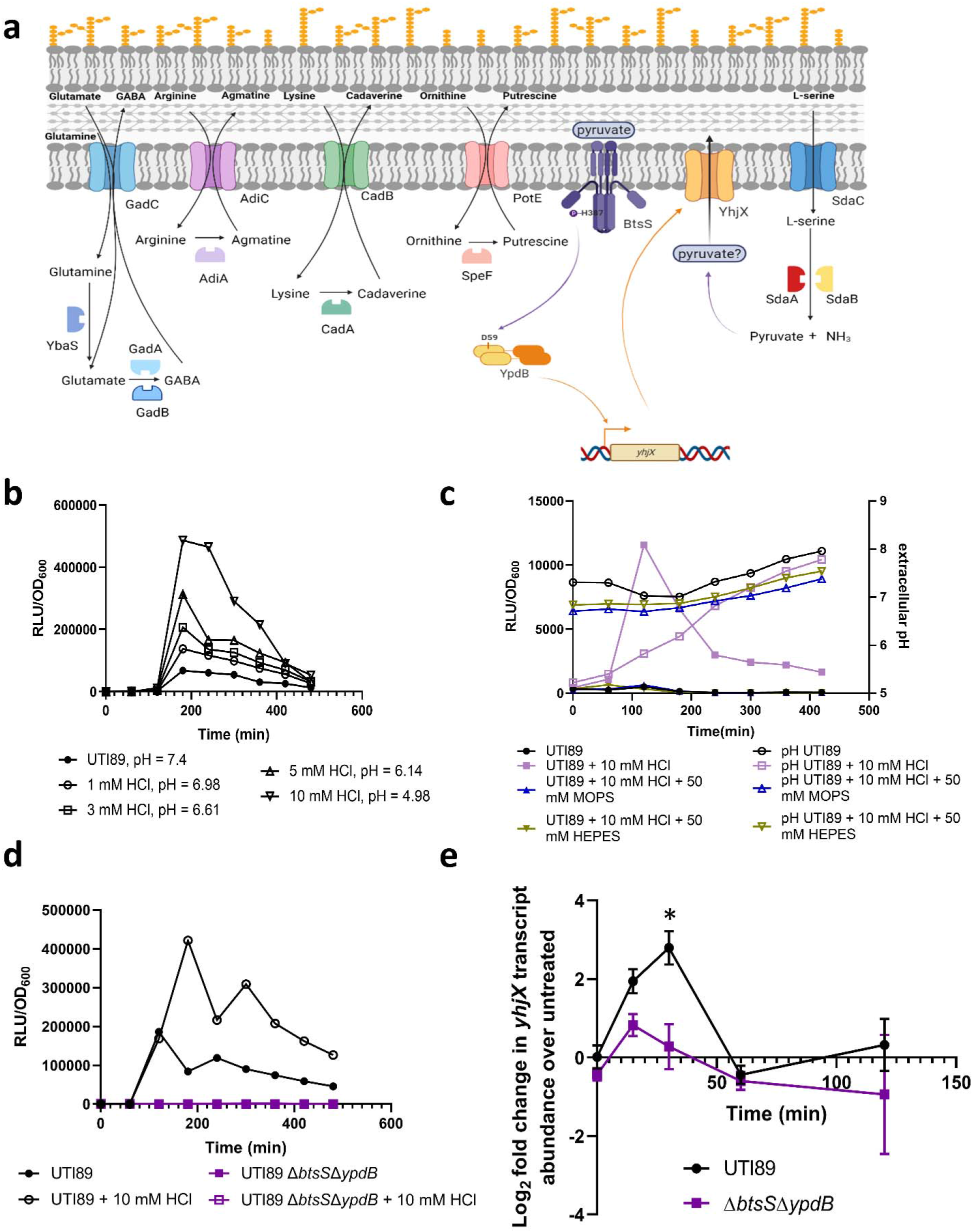
*yhjX* is upregulated in response to low pH in a manner that depends on the non-cognate two-component system BtsS and YpdB. a) Cartoon depicts currently known acid resistance mechanisms in *E. coli*, as well as a new acid resistance mechanism we propose here that depends on L-serine de-amination. Created with BioRender.com b-c) Graphs depict relative luminescence units (RLU) normalized to growth (OD_600_) over time, of UPEC strain UTI89 harboring the P*yhjX*-*lux* reporter in increasing concentration of HCl (b) or during growth in buffered media (c). d) Luciferase reporter assay of UPEC strain UTI89 the isogenic Δ*btsS*Δ*ypdB* strain grown in the presence or absence of acid. Graphs are representative of 3 independent biological repeats. E) RT-qPCR analysis of *yhjX* transcript abundance after acid stimulation in wild-type UTI89 (black) and Δ*btsS*Δ*ypdB* (purple) strains. Relative fold change was determined by the ΔΔ*C*_*T*_ method, where transcript abundances were normalized to *gyrB* housekeeping gene transcripts. \**P* = 0.0165 calculated by two-way ANOVA with Sidak’s multiple comparisons test. Error bars indicate SEM of three biological replicates.

The *yhjX* gene is under the control of a non-cognate signal transduction pair: the pyruvate sensor histidine kinase BtsS and the transcriptional regulator YpdB (24, 30). Studies have shown that pyruvate increases in the extracellular milieu as the bacterial culture senesces (23), and that this pyruvate is directly sensed by the BtsS histidine kinase, leading to subsequent upregulation of *yhjX* via the action of the YpdB response regulator (24).To determine whether acid-mediated induction of *yhjX* is dependent on BtsS and YpdB, a mutant lacking both *btsS* and *ypdB* (Δ*btsS*Δ*ypdB* (24)) was tested via our luminescent reporter assay in acidic and neutral conditions. These experiments showed no induction of *yhjX* in the Δ*btsS*Δ*ypdB* strain, regardless of pH in the culture media (Fig 1d and Supplementary file 1). To validate these findings, *yhjX* steady-state transcript over time was also monitored by RT-qPCR and TaqMan based chemistry in acidified and non-acidified cultures of wild-type UTI89 and the isogenic Δ*btsS*Δ*ypdB* mutant. Transcript abundance of *yhjX* was compared between acid stimulated and unstimulated growth conditions and normalized to the *gyrB* housekeeping gene. These analyses revealed a characteristic transcription surge (31) for *yhjX* in acidified wildtype UTI89 cultures, which was not apparent in the isogenic Δ*btsS*Δ*ypdB* (Fig 1e). To further confirm that *yhjX* is induced by low pH and not just HCl, we utilized the luminescent reporter to monitor *yhjX* induction in UTI89 and Δ*btsS*Δ*ypdB* in the presence of lactic acid, acetic acid, and pyruvic acid. All organic acids tested resulted in strong *yhjX* induction in UTI89. However, the Δ*btsS*Δ*ypdB* strain did not induce *yhjX* under any of these acidic conditions (Fig. S2 and Supplementary file 1). Together, these data confirm that *yhjX* is induced in low-pH growth conditions and that this induction depends on the presence of BtsS and YpdB.

### Induction of BtsS-YpdB in response to acid results from pyruvate produced during L-serine deamination

Previous work demonstrated that BtsS signaling is affected by L-serine levels in the growth medium (30). These studies postulated that L-serine, which is the first amino acid to be consumed by *E. coli* during growth in laboratory media (17), is converted to pyruvate by the L-serine deaminases SdaA and SdaB and then presumably exported by YhjX to serve as a positive feedback signal for BtsS ((18, 32, 33) and Fig. 1a). Given that the de-amination of L-serine also produces ammonia, which can consume a proton and thus raise intracellular pH (Fig. 1a), we asked whether the mechanism of acid stress alleviation observed in our studies depends on the import and deamination of L-serine. To test this hypothesis, we first created a series of mutants lacking the SdaC transporter (Δ*sdaC*), the SdaA or SdaB de-aminases (ΔsdaA, Δ*sdaB*) or both de-aminases (Δ*sdaA*Δ*sdaB*). The induction of *yhjX* in these mutants was tested via luciferase reporter assays either in media in which exogenous L-serine was added (Fig. 2a and Supplementary file 1), or in media acidified with HCl (Fig. 2b and Supplementary file 1). Addition of L-serine did not induce *yhjX* promoter activity in any of the *sda* mutants (Fig. 2a and Supplementary file 1). Addition of acid to the media led to decreased *yhjX* induction compared to wild-type UTI89 in all the single mutants tested and led to no *yhjX* induction in the Δ*sdaA*Δ*sdaB* strain (Fig. 2b, Supplementary file 1). Given that pyruvate is the known ligand of the BtsS sensor, our data suggests that *in-vivo* it is the de-amination of L-serine into pyruvate that actually leads to *yhjX* being induced. As further evidence that it is the generation of pyruvate driving *yhjX* induction, sodium pyruvate was added to the media of the Δ*sdaA*Δ*sdaB* strain, which caused – as expected – *yhjX* induction (Figure S3 and Supplementary file 1).

**Figure 2.**
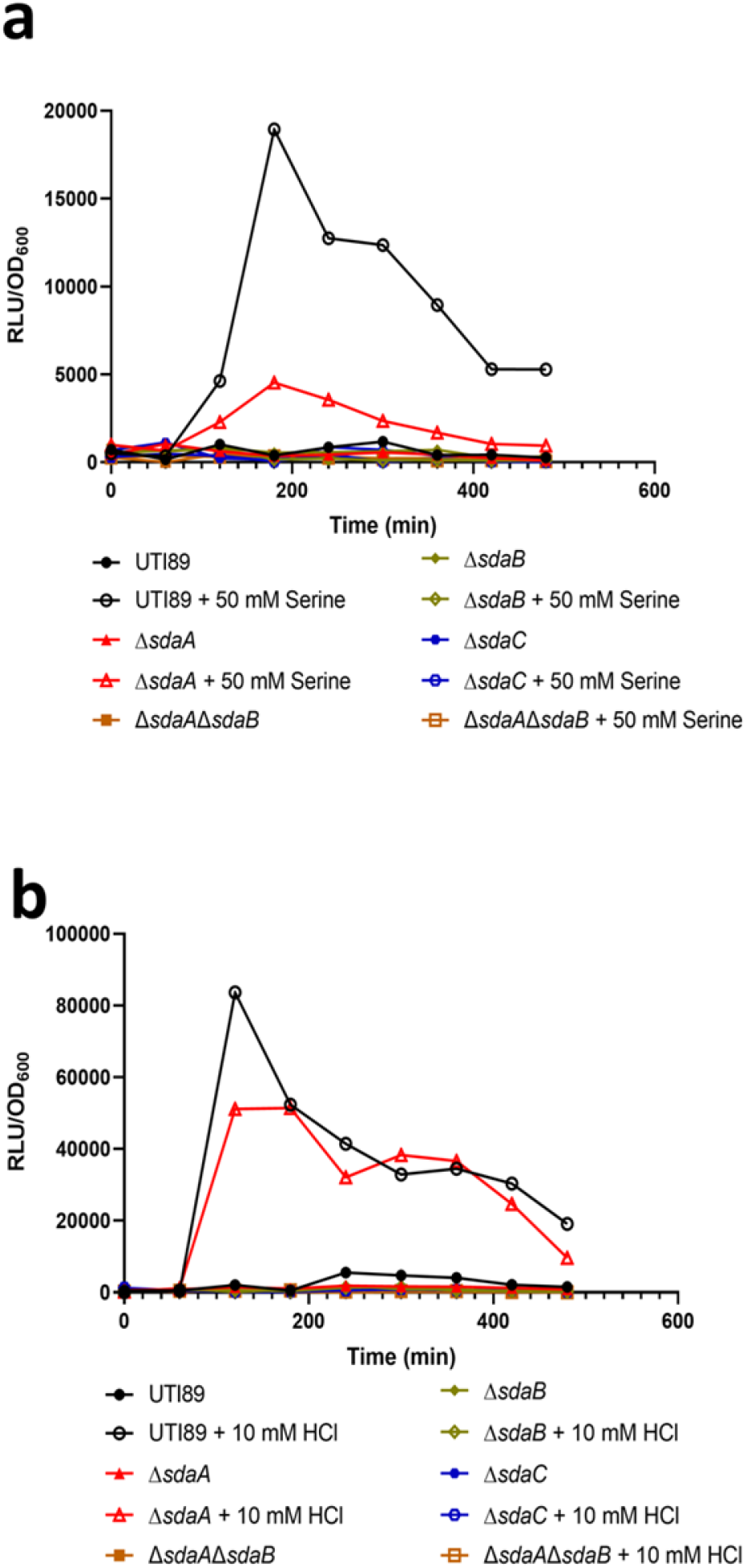
Induction of BtsS-YpdB in response to acid results from pyruvate produced during L-serine deamination. Graphs depict luciferase reporter assay performed over time of strains harboring the *yhjX* promoter reporter during growth in media supplemented with 50 mM serine (a) or 10 mM HCl (b). Deletion of serine import (*sdaC*, blue) or de-amination genes (*sdaA*, red; *sdaB*, gold) diminishes *yhjX* promoter activity. Results are representative of 3 biological repeats.

### L-serine deamination is another acid resistance mechanism in *E. coli*

If L-serine deamination is a component of the *E. coli* acid response, we reasoned that the mutant lacking the SdaA/B enzymes would display a survival defect in acidic conditions. Indeed, testing the *sda* deletion strains in acid survival assays demonstrated significant reduction in survival for all strains, compared to the wild-type (Fig. 3a). The Δ*sdaA*Δ*sdaB* strain exhibits the most pronounced acid survival defect, having an acid susceptibility profile similar to known acid resistance mutants (Fig. 3a).

**Figure 3.**
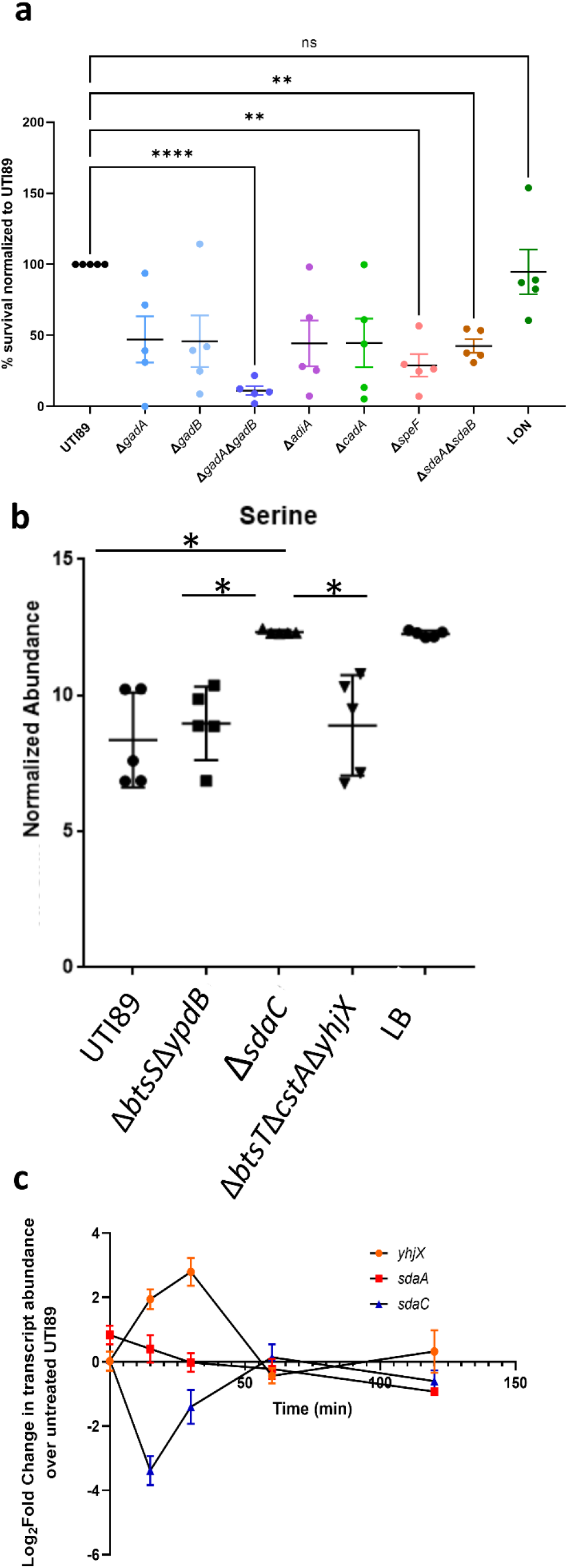
L-serine deamination is another acid resistance mechanism in *E. coli*. a) Graph depicts survival in acidic conditions, compared to the wild-type strain, of mutants deleted for decarboxylases or serine deaminases. For these assays, cultures were incubated for three hours, at which point an aliquot was collected for CFU enumeration before acid treatment. The remaining culture was treated with HCl to adjust the pH to three. Samples were incubated for an additional 30 minutes, after which they were plated for CFUs. Percent survival in acid is calculated as the number of CFUs in acid treatment, compared to untreated input control. Statistical analysis was performed by 1-way ANOVA with *post hoc* Dunnett’s multiple comparisons correction test (\*\**P*<0.005, \*\*\*\**P* <0.0001). Error bars indicate SEM of 5 biological replicates. b) UTI89, Δ*btsS*Δ*ypdB*, and Δ*sdaC* were grown until cultures reached an OD_600_ = 0.5, then 1 M HCl was added to the culture to a final concentration of 10 mM (pH=5). Cultures were incubated for another 15 minutes, then 1 ml of culture was collected. Cells were pelleted and supernatant was flash frozen and stored at -80°C prior to sample preparation. Following MS sample preparation, extracellular serine abundance was detected by LC-MS. c) qPCR analysis of *yhjX* (orange), *sdaA* (red), and *sdaC* (blue) transcript abundance after acid stimulation in wildtype UTI89. The relative fold change was determined by the ΔΔ*C*_*T*_ method where transcript abundances were normalized to *gyrB* housekeeping gene transcripts. Error bars indicate SEM of three biological replicates (\**P*< 0.005, ANOVA).

To determine if serine is imported by *E. coli* in response to a drop in pH, cell culture supernatants were analyzed for relative serine abundance by LC-MS and compared to acidified media alone as a control. Mass spectrometric measurements of serine in media alone demonstrated that baseline serine abundance level can be detected with this method (Fig. 3b and supplementary data 2). Measurements of serine in acidified media revealed a significant reduction in extracellular serine abundance in the supernatant fractions of WT UTI89, but not the Δ*sdaC* mutant (Fig. 3a and Supplementary file 2). These data indicate that serine is imported – in an SdaC-dependent manner – into the bacterial cell in response to acid stress.

Extracellular serine levels also drop in the Δ*btsS*Δ*ypdB* mutant under acidic conditions (Fig. 3b and Supplementary file 2). This observation is consistent with the notion that L-serine is imported independent of BtsS-YpdB signaling. Subsequent qPCR showed that in the wild-type strain *sdaA* and *sdaBC* transcript abundance does not change in response to acid stress (Fig. 3c), in sharp contrast to *yhjX* that displays an activation surge (Fig. 3c). Together these data demonstrate that SdaC imports L-serine in response to acidic stress and that L-serine deamination via the action of SdaA or SdaB increases *E. coli* survival under acid stress, while the pyruvate produced acts as a ligand for BtsS-YpdB signaling.

Given that L-serine de-amination leads to the production of pyruvate, which is the known ligand for BtsS (30), we hypothesized that SdaA/SdaB-mediated L-serine deamination increases intracellular pyruvate levels that could then be exported via YhjX and that this export of pyruvate may play a role in this acid stress response (Fig. 4a). To determine if extracellular pyruvate abundance changes correspond to L-serine depletion, we measured pyruvate levels in the extracellular milieu 15 minutes after acidification of the culture media, which is the same timepoint used for the serine measurements (Fig. 3). As performed for serine, cell culture supernatants were analyzed for relative pyruvate abundance by LC-MS compared to acidified media alone as a control. In wildtype UTI89, an increase in extracellular pyruvate is observed under these conditions, compared to the media control (Fig. 4b and Supplementary file 2). However, an increase in extracellular pyruvate is also observed in our Δ*btsS*Δ*ypdB* mutant strain which as demonstrated (Fig. 1) has no observable *yhjX* promoter activity (Fig. 4b and Supplementary file 2). These data suggest that the proposed export of pyruvate via YhjX is not itself a mechanism that allows *E. coli* to withstand acid stress under the conditions tested. We also measured extracellular pyruvate concentration in a UTI89 strain deficient in all known and potential pyruvate transporters (UTI89 Δ*btsT*Δ*cstA*Δ*yhjX*) (23, 34, 35). Again, we observed no significant difference in extracellular pyruvate abundance in the supernatants of these mutants, compared to the wild-type parent. This could be because pyruvate becomes protonated in the interior of the cell and freely diffuses out of the bacterial cell under the conditions tested impeding our ability to measure differences in pyruvate export. Other possibilities include the presence of a yet unidentified pyruvate transporter that exports pyruvate, or that pyruvate produced by L-serine deamination is consumed intracellularly. Whatever the reason, our data demonstrates that the import and deamination of L-serine provides acid resistance to *E. coli* but that the hypothesized exportation of pyruvate via YhjX is likely not what mediates this increased survival under acidic conditions.

**Figure 4.**
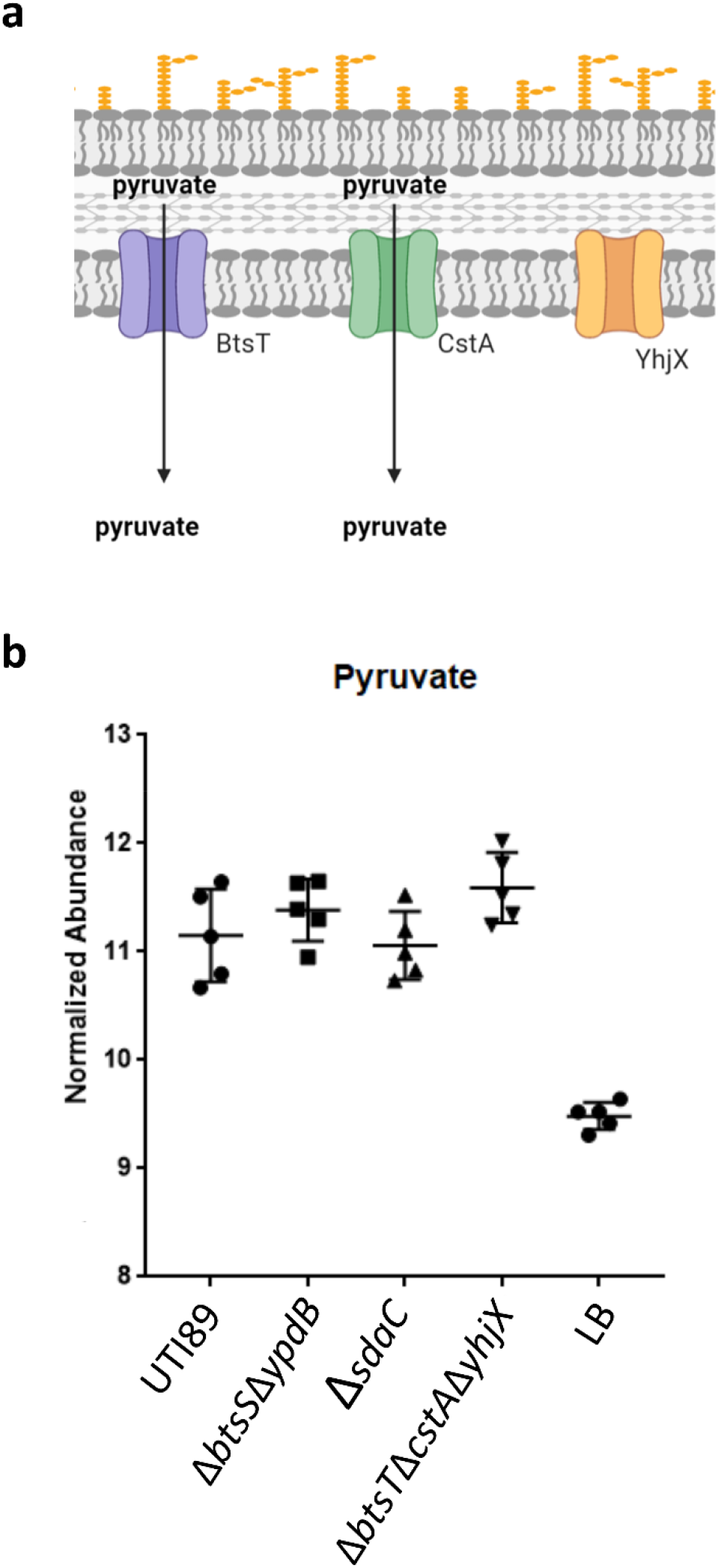
a) Cartoon depicts currently known pyruvate transporters (BtsT and CstA) and a putative pyruvate transporter (YhjX) in *E. coli*. Created with BioRender.com b) UTI89, Δ*btsS*Δ*ypdB*, Δ*sdaC*, and Δ*btsT*Δ*cstA*Δ*yhjX* strains were grown until cultures reached an OD_600_ = 0.5, then 1 M HCl was added to the culture to a final concentration of 10 mM (pH=5). Cultures were incubated for another 15 minutes, then 1 ml of culture was collected. Supernatant was collected, flash frozen, and stored at -80°C prior to sample preparation. Following MS sample preparation, extracellular pyruvate abundance was detected by LC-MS.

## Discussion

In this work, we show that L-serine deamination serves as an additional acid response mechanism in *E. coli*. Serine is the first amino acid consumed by *E. coli* when grown in complex media, despite its toxicity at higher concentrations (17). L-serine can be deaminated by the SdaA and SdaB enzymes to yield pyruvate and ammonia. *E coli* encodes multiple serine deaminase genes that function under a variety of environmental conditions (36). For instance, while both *sdaA* and *sdaB* are expressed under aerobic conditions, *sdaA* is expressed in nutrient limited minimal media, and in contrast, *sdaB* is expressed in the nutrient rich LB(19, 33, 37). The redundancy of these enzymes indicates that the deamination of serine is important for these bacteria. In this paper, we propose that L-serine importation into *E. coli* serves as an acid resistance mechanism through the production of ammonia and pyruvate.

Various bacteria are known to use ammonia to neutralize their intracellular pH, including *E. coli* (14, 38, 39). We postulate that the ammonia produced via L-serine deamination functions as a base which increases the cytoplasmic pH of *E. coli* when under acid stress. A similar role has been reported for glutamine (AR2_Q) in *E. coli* (14, 16). Deletion of the serine deaminase genes, *sdaA* and *sdaB*, results in a significant decrease in cell survival in acidic conditions compared to the wildtype strain. This decrease in cell survival is comparable to that observed in other acid resistance mechanism mutants of *E. coli*. All together, these data suggest that L-serine deamination serves as another acid resistance mechanism in *E. coli*.

Serine is taken up from the media in the transition from logarithmic phase to stationary phase (17, 33). This timing of serine consumption coincides with the activation of BtsS-YpdB signaling (Fig 1b-d). The onset of stationary phase is concomitant with the induction of many stress response genes, including those involved in AR2 (40, 41). The timing of serine import may be indicative of its role in alleviating stress in *E. coli* during the transition to stationary phase of growth, as nutrients are depleted from the media. This is also the phase of growth cells are in when they enter the host (15). Loss of serine deaminase activity was previously shown to result in changes to cell shape due to interference with cell wall synthesis (19, 36). These serine deaminase mutants exhibit increased filamentation. In the context of UTI, filamentous UPEC are deficient in invading bladder cells, forming secondary intracellular bacterial communities, and in establishing quiescent intracellular reservoirs (42, 43). Thus, loss of serine deamination would detrimentally affect *E. coli’s* ability to form reservoirs in addition to its ability to tolerate low pH. Evolution of mechanisms that utilize various amino acids for acid resistance would allow bacteria to seamlessly adapt to different host niches with differing resource availabilities during the infection process.

It has been shown that up to 51% of serine flux is directed to the production of pyruvate (17), which can presumably be shunted into the TCA cycle for energy production. Here we demonstrate a connection between serine deamination and the pyruvate responsive cross regulating two-component system, BtsS-YpdB. BtsS-YpdB activation occurs in response to increased pyruvate and leads to the upregulation of *yhjX* transcription. Loss of L-serine deaminases result in ablation of *yhjX* promoter induction in response to serine, indicating that pyruvate produced via serine deamination induces activation of the BtsS-YpdB system. Our metabolomics analysis demonstrates that extracellular serine levels drop in the Δ*btsS*Δ*ypdB* mutant under acidic conditions indicating that BtsS-YpdB activation occurs after L-serine is imported. In this work we also show that *yhjX* is also induced by the BtsS-YpdB system in response to acidic pH (Fig 1b-d). Although this activation in response to low pH was dose dependent, considering the data that suggests *yhjX* is induced through serine deamination and that pyruvate is a known ligand for BtsS, *yhjX* activation in response to low pH appears to be a response to the pyruvate produced by serine deaminases. The function of YhjX remains elusive. We, and others, have hypothesized that YhjX could serve as a pyruvate transporter. Our metabolomics experiments indicate that this may not be the case. When comparing wildtype UPEC to the isogenic deletion strains lacking the known pyruvate transporters and YhjX, there was no difference in extracellular pyruvate concentrations (Fig 4b). Arguably, these data were collected at a single time point. It is possible that extending the time course of sampling could show differences in pyruvate excretion by the tested strains. It is also possible that YhjX is critical because it functions as the transporter of a different metabolite that is involved in acid resistance under the conditions tested. Recent studies have indicated that pyruvate – or metabolites downstream of pyruvate utilization pathways – play a role in increasing acid resistance in *E. coli* (44). It is possible that pyruvate sensing through BtsS-YpdB and upregulation of YhjX is involved in downstream utilization of pyruvate during the acid response.

In summary we show that in acidified media, cells scavenge L-serine from the growth media via the SdaC transporter and de-aminate it via the action of SdaA and SdaB. Deletion of both deaminase genes renders *E. coli* susceptible to acid stress similarly to known acid stress deletion mutants. We therefore propose that the importation and deamination of serine represents a previously uncharacterized acid response mechanism in *E. coli*.

## ACKNOWLEDGEMENTS

We thank members of the Hadjifrangiskou and Schmitz labs for helpful discussions and critical reading of the manuscript. This work was supported by the National Institutes of Health under the following grants: P20DK123967 (MH), T32 GM007569 (JRB) and 2T32AI112541-06 (MAW). This work was supported in part using the resources of the Center for Innovative Technology at Vanderbilt University.

## AUTHOR CONTRIBUTIONS

MAW, BDS and MH conceived the study, performed the experiments, analyzed the data, and composed the manuscript. AB and JRB performed experiments, analyzed data, and edited the manuscript. ACR, SGC, SDS and JAM optimized mass spectrometry methods, executed the metabolomics experiments, performed metabolomics data analysis, edited and reviewed the manuscript.

